# A Large-scale Drug Repositioning Survey for SARS-CoV-2 Antivirals

**DOI:** 10.1101/2020.04.16.044016

**Authors:** Laura Riva, Shuofeng Yuan, Xin Yin, Laura Martin-Sancho, Naoko Matsunaga, Sebastian Burgstaller-Muehlbacher, Lars Pache, Paul P. De Jesus, Mitchell V. Hull, Max Chang, Jasper Fuk-Woo Chan, Jianli Cao, Vincent Kwok-Man Poon, Kristina Herbert, Tu-Trinh Nguyen, Yuan Pu, Courtney Nguyen, Andrey Rubanov, Luis Martinez-Sobrido, Wen-Chun Liu, Lisa Miorin, Kris M. White, Jeffrey R. Johnson, Christopher Benner, Ren Sun, Peter G. Schultz, Andrew Su, Adolfo Garcia-Sastre, Arnab K. Chatterjee, Kwok-Yung Yuen, Sumit K. Chanda

## Abstract

The emergence of novel SARS coronavirus 2 (SARS-CoV-2) in 2019 has triggered an ongoing global pandemic of severe pneumonia-like disease designated as coronavirus disease 2019 (COVID-19). To date, more than 2.1 million confirmed cases and 139,500 deaths have been reported worldwide, and there are currently no medical countermeasures available to prevent or treat the disease. As the development of a vaccine could require at least 12-18 months, and the typical timeline from hit finding to drug registration of an antiviral is >10 years, repositioning of known drugs can significantly accelerate the development and deployment of therapies for COVID-19. To identify therapeutics that can be repurposed as SARS-CoV-2 antivirals, we profiled a library of known drugs encompassing approximately 12,000 clinical-stage or FDA-approved small molecules. Here, we report the identification of 30 known drugs that inhibit viral replication. Of these, six were characterized for cellular dose-activity relationships, and showed effective concentrations likely to be commensurate with therapeutic doses in patients. These include the PIKfyve kinase inhibitor Apilimod, cysteine protease inhibitors MDL-28170, Z LVG CHN2, VBY-825, and ONO 5334, and the CCR1 antagonist MLN-3897. Since many of these molecules have advanced into the clinic, the known pharmacological and human safety profiles of these compounds will accelerate their preclinical and clinical evaluation for COVID-19 treatment.

## Introduction

In December 2019, the novel SARS coronavirus 2 (SARS-CoV-2) was identified as the causative agent of a severe pneumonia-like coronavirus disease (COVID-19) outbreak in Wuhan in the Hubei province of China^1^. SARS-CoV-2 is an enveloped, positive-sense, single-stranded RNA betacoronavirus, related to the viruses that caused severe acute respiratory syndrome (SARS) and Middle East respiratory syndrome (MERS) in 2002-2004 and 2012-present, respectively. The World Health Organization (WHO) declared the rapidly spreading disease a pandemic on March 11^th^, 2020, and, as of April 16^th^, more than 2.1 million confirmed cases and 139,500 deaths have been recorded worldwide in 213 countries^2^. The WHO estimates the global case fatality rate (CFR) at 3.4% of those infected, though the number of actual infections is likely much higher than the number of reported cases^3,4^. Typical COVID-19 symptoms include fever, cough, anosmia, headache, anorexia, myalgia, and, in the most severe cases, viral-induced pneumonia accompanied by prolonged and systemic cytokine release^5,6^. Notably, the levels of IL-6 have been reported to correlate with respiratory failure, and inhibitors are currently being pursued in clinical studies for the amelioration of virus-induced inflammatory responses^7^. Patients with pre-existing chronic conditions such as hypertension, diabetes, and asthma, as well as those 65 years or older, are at a higher risk of severe disease outcome^8^. The underlying basis for these differential outcomes is yet unknown. Together, the still accelerating rate of community transmission and severity of the symptoms have placed an unprecedented burden on the medical supply chain and health care system in Italy, Spain, and the U.S.^9-11^, with similar scenarios playing out or anticipated in other countries.

While the FDA has recently granted the antimalarial drug hydroxychloroquine sulfate (also known as hydroxychloroquine) emergency use authorization (EUA) for COVID-19 treatment, at present, there is no vaccine or approved antiviral therapeutic agent available^12^. Thus, there is an urgent and critical need to identify novel medical countermeasures both for prophylactic and treatment use. Since the production of a vaccine could take 12-18 months^13^, and *de novo* development of therapies usually requires 10-17 years^14^, repositioning clinically evaluated drugs represents one of the most practicable strategies for the rapid identification and deployment of treatments for emerging infectious diseases such as COVID-19.

Toward this end, in addition to many anti-immune treatments not addressed in this paper, many investigational clinical trials using repurposed drugs for evaluation of direct antiviral activity have already been launched. Those include multiple antiviral and antimalarial medicines. Early results of a multicenter trial in China suggested that the antimalarial drug, chloroquine, may limit exacerbation of pneumonia and shorten viral replication and course of disease^15^. A French study that used hydroxychloroquine together with azithromycin reported a significant reduction in viral load in COVID-19 patients when used in combination^16^. However, a sufficiently powered case-control study has not yet been reported, and thus it is unclear if there are therapeutic benefits of chloroquine administration to SARS-CoV-2-infected patients, although several concerns are being raised recently, due to the severe cardiac complications potentially resulting from the use of this treatment in COVID-19 patients^17,18^.

The repurposing of several approved antiviral therapies have all been the focus of clinical investigations, including HIV-1 protease inhibitors lopinavir/ritonavir (Kaletra, Aluvia by AbbVie)^19^, hepatitis C virus protease inhibitor danoprevir (Ganovo, Ascletis Pharma)^20^, and the influenza antiviral favipiravir (T-705, Avigan)^21^. Most notably, ten clinical trials at more than 50 global sites are underway to investigate remdesivir (GS-5734), an investigational antiviral originally developed by Gilead Sciences to treat Ebola virus infection^22^. Remdesivir, an adenosine analogue, is a viral RNA polymerase inhibitor that causes premature termination of transcription when incorporated into nascent viral RNA^23^. The drug has demonstrated *in vitro* and *in vivo* activity in animal models against both MERS and SARS^24,25^, as well as potent antiviral activity in Vero E6 against a clinical isolate of SARS-CoV-2^26^. Pending results of several randomized (n = 308) clinical trials are expected to provide definitive insight into the efficacy of remdesivir as a therapeutic solution for the treatment of COVID-19. However, a well-powered randomized controlled trial has yet to demonstrate definitive evidence of antiviral efficacy for remdesivir or any other potential therapeutic.

While these targeted repurposing strategies provide potentially rapid trajectories toward an approved treatment, an unbiased large-scale evaluation of known drugs and clinical candidates can identify additional unanticipated therapeutic options with accelerated evaluation for the treatment of COVID-19 disease. Here, we describe a high-throughput repositioning screen using the commercial library of 1,280 pharmacologically active compounds LOPAC®1280 and the ReFRAME (<u>Re</u>purposing, <u>F</u>ocused <u>R</u>escue, and <u>A</u>ccelerated <u>Me</u>dchem) drug collection, a comprehensive open-access library of ∼12,000 clinical-staged or approved small molecules respectively, to identify existing drugs that harbor antiviral activity against SARS-CoV-2 in a cell-based assay^27,28^. The ReFRAME library has previously been used to successfully identify the anti-inflammatory auranofin as a potential therapy for tuberculosis^29^, the approved drug clofazimine as a potent antiparasitic compound that is now being tested for efficacy against *Cryptosporidium*^30^, and has also been used to identify and optimize the FDA-approved antifungal drug itraconazole as a novel efficacious molecule suitable for chronic administration as an anti-fibrotic^31^. Importantly, the ReFRAME library is unique in that it is the only repurposing collection we are aware of where nearly 50% of the library was derived from custom synthesis, as commercially available sources of these clinical molecules were not available^32^. Each of the molecules in this collection has been previously optimized for efficacy, safety, and bioavailability. Therefore, this enables leveraging of the considerable investment in research and development to compress the timeline required for drug discovery and development^33^. For example, repositioning is estimated to reduce the typical 10-17 year development process to 3-12 years^34^, and under emergency use authorization (EUA) for chemical, biological, radiological, and nuclear (CBRN) threats during public health emergencies, such as the SARS-CoV-2 pandemic, this may be abbreviated to a <6-month time frame.

Here, we describe a high-throughput analysis of the ReFRAME library to identify inhibitors of SARS-CoV-2 replication in mammalian cells, and identified several targets and mechanistic classes that were highly enriched, including aldose reductase inhibitors, retinoic acid receptor antagonists, benzodiazepine receptor agonists, regulators of cholesterol homeostasis and antimalarial compounds. Validation studies further confirmed 30 known drugs to inhibit viral replication, including four molecules previously approved by the FDA (clofazimine, acitretin, tretinoin, and astemizole) or registered outside the US (tamibarotene). Dose response studies have thus far characterized 7 compounds that exhibit a range of effective concentrations (EC_50_) that are consistent with potential clinical efficacy. These include a PIKfyve kinase inhibitor that has reached Phase II clinical trials (Apilimod), and cysteine protease inhibitors (MDL-28170, Z LVG CHN2, VBY-825, and ONO 5334) that are in various phases of preclinical and clinical development. In addition, the preclinical ion channel blocker AMG-2674 and the ion channer blocker and antimalarial drug hanfangchin A, the phase I proton pump inhibitor YH-1238, as well as the G-protein receptor antagonists MLN-3897 and SDZ-62-434^35^, which are in phase II and phase I clinical evaluation respectively, were also found to possess antiviral activity against SARS-CoV-2. Rapid experimental and clinical evaluation of these therapeutics for *in vivo* antiviral efficacy and amelioration of disease-associated pathologies can provide an important opportunity for the accelerated development of potential therapies for COVID-19 treatment.

## Materials and Methods

### Cells and Viruses

SARS-CoV-2 HKU-001a strain was isolated from the nasopharyngeal aspirate specimen of a laboratory-confirmed COVID-19 patient in Hong Kong^36^. SARS-CoV-2 USA-WA1/2020 strain, isolated from an oropharyngeal swab from a patient with a respiratory illness who developed clinical disease (COVID-19) in January 2020 in Washington, USA, was obtained from BEI Resources (NR-52281). The virus was propagated in Vero E6 (ATCC® CRL-1586™) cells transfected with exogenous human ACE2 and TMPRSS2 and stored at −80 °C in aliquots. Plaque forming unit (PFU) and TCID_50_ assays were performed to titrate the cultured virus. Vero E6 and Huh-7 cells (Apath LLC, Brooklyn) were maintained in Dulbecco’s modified eagle medium (DMEM, Gibco) supplemented with 10 % heat-inactivated fetal bovine serum (FBS, Gibco), 50 U/mL penicillin, 50 µg/mL streptomycin, 1 mM sodium pyruvate (Gibco), 10 mM HEPES (Gibco), and 1X MEM non-essential amino acids solution (Gibco). Huh-7 cells were transfected with PLVX-ACE2 and PLX304-TMPRSS2 prior to infection. All experiments involving live SARS-CoV-2 followed the approved standard operating procedures of the Biosafety Level 3 facility at the University of Hong Kong^37^ and Sanford Burnham Prebys Medical Discovery Institute.

### Chemical libraries

The LOPAC®1280 library is a collection of 1,280 pharmacologically active compounds, covering all the major target classes, including kinases, GPCRs, neurotransmission and gene regulation (Sigma). The ReFRAME (<u>Re</u>purposing, <u>F</u>ocused <u>R</u>escue, and <u>A</u>ccelerated <u>Me</u>dchem) library, built at the California Institute for Biomedical Research (Calibr)^27^, contains approximately 12,000 high-value molecules assembled by combining three databases (Clarivate Integrity, GVK Excelra GoStar and Citeline Pharmaprojects) for fast-track drug discovery. This library contains US Food and Drug Administration (FDA)-approved/registered drugs (∼35 %), investigational new drugs (∼58 %), and preclinical compounds (∼3 %).

### Drug screening

Compounds from the LOPAC®1280 and ReFRAME library were transferred into F-BOTTOM, µCLEAR®, BLACK 384-well plates (Greiner) using an Echo 550 Liquid Handler (Labcyte). All compounds were diluted in culture media to a final concentration of 5 µM during screening. Briefly, Vero E6 cells were seeded in 384-well plates, on top of pre-spotted compounds, at a density of 3,000 cells per well in 40 µl using a microFlo™ select dispenser (BioTek Instruments). Sixteen hours post-seeding, the cells were infected by adding 10 µl of SARS-CoV-2 per well at an MOI of 0.01. Cytopathic effect (CPE) was indirectly quantified as the presence of ATP in live cells by using the CellTiter-Glo (Promega) luminescent cell viability assay at 72 hours post-infection. Data were normalized to the median of each plate. For the ReFRAME library, the Z-score was calculated based on the log2FC with the average and standard deviation of each plate. The screen was performed in duplicate by running the assay in parallel for the LOPAC®1280 library or as two independent experiments for the ReFRAME collection. Twenty-eight compounds from the LOPAC®1280 were selected according to the cutoff of >5*Stdev Log2FC and included in a dose-response confirmation assay. Compounds from the ReFRAME collection were ranked according to their Z-score. The top 100 hits from each replicate were selected (25 overlapping). Seventy-five additional hits were chosen according to their ranking based on the average Z-score. The last 48 hits were selected according to drug target and pathway enrichment analysis. The 298 prioritized hits were included in a dose-response confirmation assay.

### Dose Response Curves, EC_50_ Calculations, and Orthogonal Validation

The selected hits were further validated by immunofluorescence in an 8-point dose response experiment to determine EC_50_ and CC_50_ through a cell-based high-content imaging assay, labeling the viral nucleoproteins within infected cells. Three thousand Vero E6 cells were added into 384-well plates pre-spotted with compounds, in a volume of 40 µl. The final concentration of compound ranged from 1.1 nM to 2.5 µM. Sixteen hours post-seeding, 10 µl of SARS-CoV-2 USA-WA1/2020 were added to each well, at an MOI of 0.75. Twenty-four hours post-infection, cells were fixed with 5 % paraformaldehyde for 4 hours and permeabilized with 0.5 % Triton X-100 for 5 min. After blocking with 3 % bovine serum albumin (BSA) for 30 min, the cells were incubated for 1 hour at room temperature with rabbit-anti-SARS-CoV-1 nucleoprotein serum, which exhibits strong cross-reactivity with SARS-CoV-2. After two washes with phosphate-buffered saline (PBS), the cells were incubated with Alexa Fluor 488-conjugated goat-anti-rabbit IgG (Thermo Fisher Scientific, USA) for 1 hour at room temperature. After two additional washes, PBS supplemented with 0.1 µg/ml antifade-4 6-diamidino-2-phenylindole (DAPI) (BioLegend, USA) was added to the cells for at least 30 min before imaging. Images were acquired using the Celigo Image Cytometer (Nexcelom). The assay results and data analysis enabled us to determine infectivity and viability/cytotoxicity. Based on all infectivity and cytotoxicity values, a 4-parameter logistic non-linear regression model was used to calculate EC_50_ and CC_50_ concentration values.

### Enrichment analysis

Compounds were annotated in the three databases used to assemble the ReFRAME library (Clarivate Integrity, GVK Excelra GoStar and Citeline Pharmaprojects) according to a variety of properties, including targets, pathways, indications, and mechanisms of actions (MOA). Each annotation property was tested for enrichment among the screening hits using the GSEA software^38,39^. The compounds annotated for each property were treated as a “gene set”. For each set of vendor annotations, the background compound set was defined as the set of compounds annotated for any property by that vendor. Enrichment results at p < 0.05 and FDR q-value < 0.25 were defined as significant. Additional enrichment analyses were performed using the free online meta-analysis tool Metascape^40^.

### Expression analysis

Gene expression analysis was conducted using single-cell RNA profiling data of samples from four macro-anatomical locations of human airway epithelium in healthy living volunteers^41^. For each gene, the fraction of cells with non-zero expression values was calculated in nasal, tracheal, intermediate, and distal samples from multiple donors. Values for each sampling location were averaged across donors. To analyze gene expression levels in different cell types, the fractions of cells with non-zero expression values were determined in all cells of a given cell type across samples. Cell types with a total of less than 250 cells detected were excluded from analysis. Clustered heat maps were generated in R using the pheatmap and viridis packages.

## Results

### Optimization of a high-throughput screen for inhibitors of SARS-CoV-2 Replication

One of the most efficient ways to identify antiviral candidates against an emergent virus, such as SARS-CoV-2, that can be rapidly evaluated in clinical trials is to repurpose clinically assessed drugs. Given the urgent need for therapeutics to treat SARS-CoV-2 infection, we developed a high-throughput assay to screen a comprehensive repurposing library. Vero E6 cells, kidney epithelial cells derived from an African green monkey, have been shown to be highly permissive to SARS-CoV-2 infection^42^ and viral replication can be assessed through measurement of viral-induced cytopathic effects (CPE)^43^. A clinical isolate of the SARS-CoV-2 virus (SARS-CoV-2 HKU-001a)^36^ was utilized for assay development and screening. The assay parameters, including cell seeding density, multiplicity of infection (MOI), and timepoints, were optimized in Vero E6 cells by measuring virus-induced CPE using CellTiter-glo, which quantifies cellular ATP levels, in a 384-well format. Maximal dynamic range and reproducibility were found at conditions of 3,000 cells/well, infection at an MOI of 0.01, and CPE measurement at 72 hours post-infection (**Figure 1A**; data not shown).

**Figure 1.**
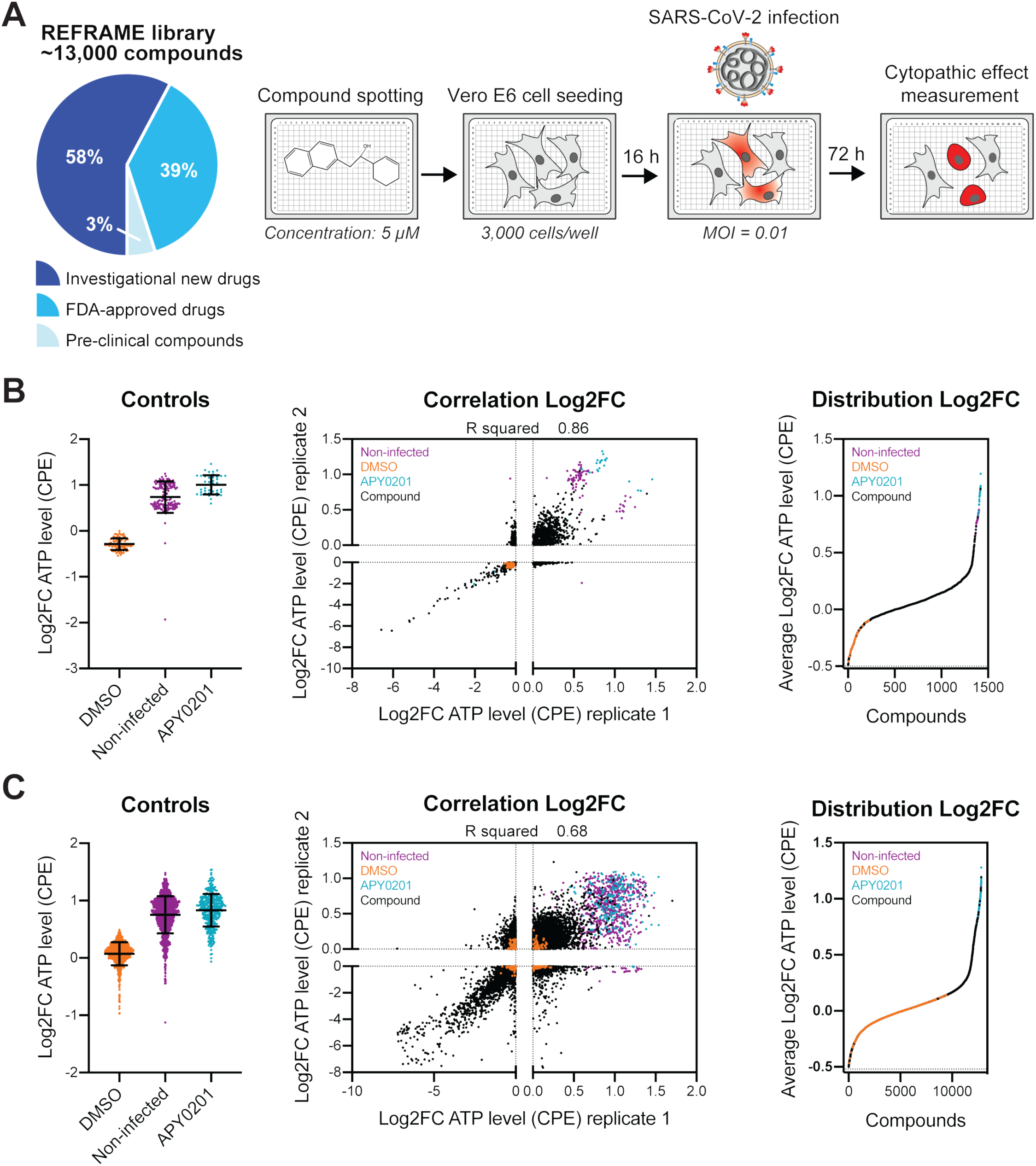
A) Screen schematic. Distribution of the approximately 12,000 compounds in the ReFRAME collection across different stages of clinical development. Compounds were pre-spotted in 384-well plates at a final concentration of 5 µM. 3,000 Vero E6 cells were added to each well and pre-incubated with each compound for 16 h, followed by infection with a clinical isolate of SARS-CoV-2 (HKU-001a) at an MOI of 0.01. ATP levels in each well were measured 72 h post-infection by the Cell Titer Glo viability assay, as a readout of the cytopathic effect (CPE) induced by the virus. B) LOPAC®1280 library primary screen. The left graph shows the Log2 fold change (Log2FC) of ATP levels after normalization to the median of each plate for all positive (APY0201) and negative (DMSO) controls as well as for non-infected cells, across all screening plates. The correlation plot in the middle panel indicates the Log2FC of each compound in the two replicates. The distribution of each compound according to the average of the log2C of each replicate (right panel) is also represented. Each dot indicates the log2FC of each drug in each replicate of the screen (black dots). Values corresponding to DMSO (orange dots), APY0201 (cyan dots) and non-infected cells (purple dots) are also represented. R squared indicates the correlation coefficient for the replicates. C) ReFRAME collection primary screen. The left graph represents the Log2 fold change (Log2FC) of ATP levels after normalization to the median of each plate for all positive (APY0201) and negative (DMSO) controls as well as for non-infected cells, across all screening plates. The correlation plot in the middle panel indicates the Log2FC of each compound in the two replicates. The distribution of each compound according to the average of the log2C of each replicate (right panel) is also represented. Each dot indicates the log2FC of each drug in each replicate of the screen (black dots). Values corresponding to DMSO (orange dots), APY0201 (cyan dots) and non-infected cells (purple dots) are also represented. R squared indicates the correlation coefficient for the replicates.

In an effort to assess robustness and reproducibility of the optimized assay in a high-throughput screening (HTS) configuration, we initially evaluated the assay utilizing the collection of 1,280 known bioactive molecules LOPAC®1280. Upon initiation of the screening effort, a compound with activity against SARS-CoV-2 in Vero E6 cells had not been reported. Based on studies that indicate that inhibition of the PIKfyve kinase inhibits low pH-dependent entry of viruses such as Ebola^44,45^, we evaluated the potential antiviral activity of the PIKfyve kinase inhibitor APY0201 against SARS-CoV-2. Compared to vehicle (DMSO), cells dosed with 1 µM APY0201 harbored a 2.5X increase in cell viability, reflecting reduced CPE after viral challenge, which was comparable to the non-infected control (**Figure 1B, left panel**). These data confirmed the antiviral activity of APY0201 against SARS-CoV-2 and enabled us to establish a reliable dynamic range based on the activity of a positive control. Vero E6 cells were seeded in 384-well plates with pre-spotted compounds from the LOPAC®1280 library at a 5 µM (final) concentration. After 16 hours, cells were infected with SARS-CoV-2 (MOI = 0.01) in the presence of compound, and at 72 hours post-infection CPE induced by the virus was quantified through previously described measurement of cell viability. Duplicates of each plate were run in parallel and the value corresponding to each well was normalized to the median of each plate and used to calculate the log base 2 of the fold change (Log2FC). Based on the activity of APY0201, the average Z’ factor for the 5 plates in duplicate was 0.4, and the correlation coefficient for the duplicates (R^2^) was 0.81 (**Figure 1B**), although an usual distribution of the positive controls in the correlation plot was observed. Twenty-eight compounds were selected for further study based on activities in both screens. These included the HIV protease inhibitor nelfinavir mesylate hydrate, the calpain and cathepsin B inhibitor MDL28170 and the antagonist of the serotonin receptors 5-HT1B and 5-HT1D GR 127935 hydrochloride hydrate, which have all been shown to efficiently block either SARS-CoV-1 or 2 infection *in vitro* or *in vivo*^46-50^, thus assessing the sensitivity of our screening conditions in identifying compounds with known antiviral activity (**Figure 1B, top right middle panel)**.

### Repositioning analysis of the ReFRAME Drug Repurposing Library

Since the results of the LOPAC®1280 HTS analysis indicated that these assay conditions were suitable to progress to a large-scale screen, we used this experimental design to screen the comprehensive ReFRAME drug repurposing collection. This library is an inclusive collection of nearly 12,000 chemical compounds, that have been either FDA-approved or registered outside the US, entered clinical trials, or undergone significant pre-clinical characterization^27^. Specifically, 11,987 compounds were arrayed in 384-well plates at a final concentration of 5 µM. As with the previous assay, Vero E6 cells were seeded into each well pre-spotted with compound, infected 16 h later with SARS-CoV-2 (MOI = 0.01) and at 72 hours post-infection, CPE was determined. Analysis of the average Z’ factor calculated on the activity of APY0201 was determined to be 0.51, reflecting an acceptable assay dynamic range (**Figure 1C, left panel**). The screen was then repeated as an independent replicate and the correlation coefficient (R^2^) for the two replicates was determined to be 0.68 (**Figure 1C, middle panel**). Data were normalized to the median of each plate and used to calculate the Log2FC. The distribution of the compounds based on the average of their Log2FC calculated within the replicates is shown in **Figure 1C** (**right panel**). Z-scores were then calculated per plate, based on the Log2FC values (**Figure S1**).

To elucidate targets, pathways, indications, and mechanisms of actions (MOA) enriched among hits in the primary screen, compounds in the collection were classified based on their reported target annotation. Gene Set Enrichment Analysis (GSEA) was used to assess the distribution of molecules with similar targets, functional categories or MOA across the screen^51,52^. We next examined whether any target, pathway, indication, or MOAs was enriched across the entire dataset. Compounds in the collection were classified based on their reported annotations from three commercial vendors, and each annotation group was tested for enrichment among hits using the GSEA software^38,39^. Based on a nominal P-value cutoff of 0.05 and FDR q-value < 0.25, we found that 15 target sets were enriched in our ranked hit list (**Figure 2A and Figure S2**). These enriched targets and biological processes include allosteric modulators of the benzodiazepine and retinoic receptors, cytosolic NADPH-dependent oxidoreductase aldose reductase, potassium channels, cholesterol homeostasis, and serine proteases. Antimalarials including chloroquine derivatives (i.e. Amopyroquine and AQ-13) were also enriched, although the FDR q-value was below the established threshold (0.33) (**Figure 2A; Figure S2 and Table S2**).

**Figure 2.**
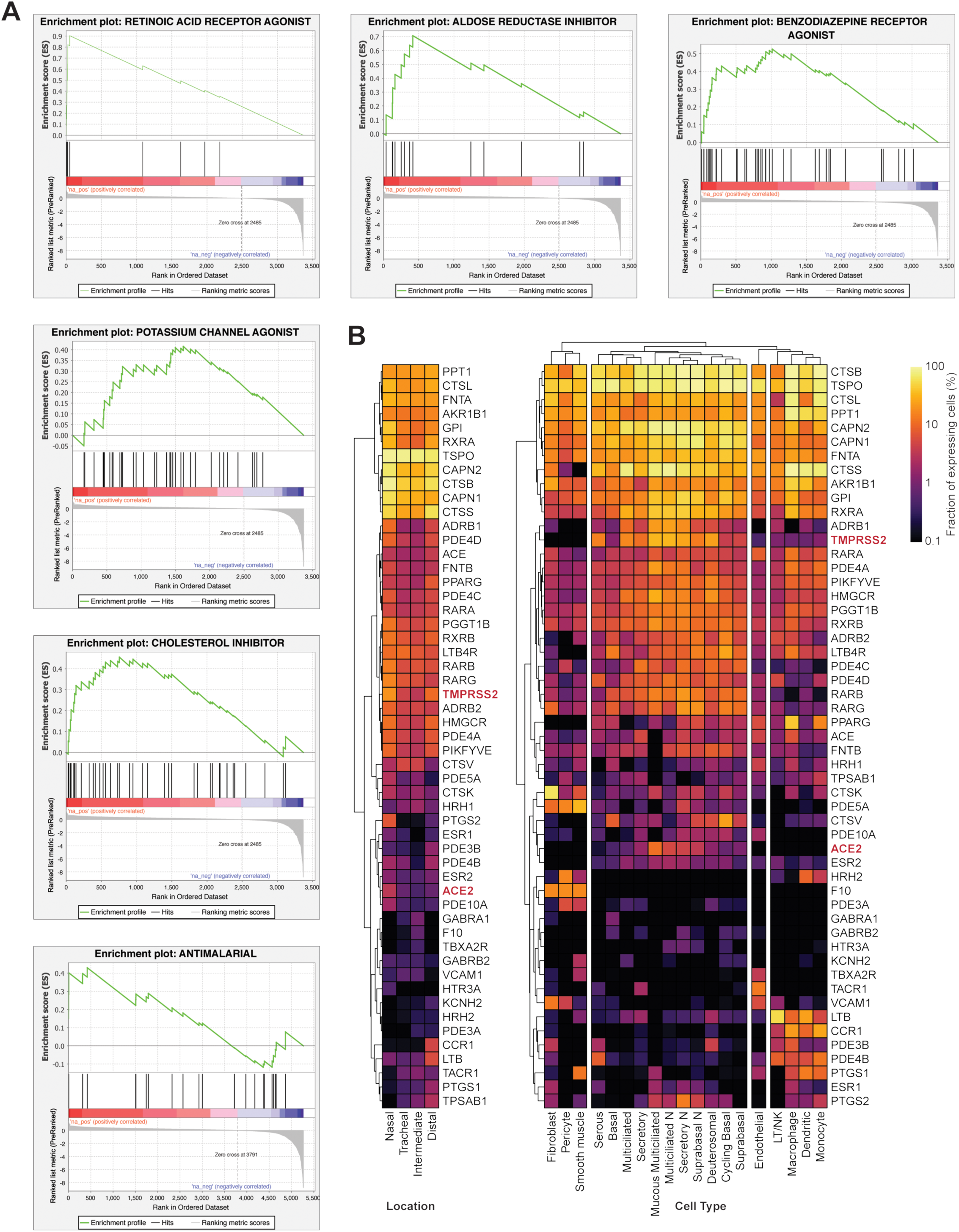
A) Gene set enrichment analysis (GSEA) of primary screening data according to the average Z′ factor. GSEA enrichment plots of five target clusters that are enriched were represented including retinoic acid receptor agonist, benzodiazepine receptor inhibitor, aldose reductase agonist, potassium channel agonist, cholesterol inhibitor, and antimalarial (P-value < 0.05, FDR q-value < 0.25). Antimalarials, enriched althought not significantly (FDR q-value 0.33), were also included as relevant class. B) Expression of ACE2, TMPRSS2, and select target genes was analyzed using single-cell RNA profiling data from human airway samples of healthy donors. Clustered heat maps show the fraction of gene-expressing cells separated by sampling locations (left panel) or cell type (right panel).

SARS-CoV-2 primarily infects the epithelial cells in the respiratory tract^53^. To elucidate the expression pattern of the genes that have been annotated to be targeted by putative antiviral compounds, we compared the expression of drug target genes enriched in the compound screen across cell types within the respiratory tract^41^. Although both the entry receptor for SARS-CoV-2, ACE2, and the priming protease TMPRSS2, were found to be expressed within specific anatomical sampling locations in the respiratory tract (**Figure 2B, left heatmap**), ACE2 expression was found to be restricted to epithelial cell types including multiciliated, nasal, deuterosomal, secretory, and basal cells (**Figure 2B, right heatmap**). Nonetheless, ACE2 expression has been reported to be induced by type-I interferon^90^. Notably, a majority of the mapped targets of active compounds also harbored expression in relevant respiratory epithelial cells, suggesting these may be physiologically relevant drug targets (**Figure 2B**). Further pathway analyses of these enriched MOAs and targets revealed enrichment in genes involved in nuclear receptor pathways, GPCR ligand binding and signaling, and calcium signaling (**Figure S3**), underscoring the potential critical role of these molecular circuits in cellular control of the SARS-CoV-2 life cycle^40^.

### Orthogonal validation of selected anti-SARS-CoV-2 compounds

To select candidates for validation studies, compounds were ranked according to their Z-score in the primary screen (Figure S1), and 100 compounds from replicate 1 were prioritized based on this ranking, and an additional 75 molecules were selected that were only found in replicate 2. 75 compounds were also selected according to their ranking calculated based on the average Z-score between the replicates, and an additional 48 hits were chosen because they were classified within an enriched drug target and pathway class (see above).

We initially assessed the activity of hits at half the original screening concentration (2.5 µM) using an orthogonal assay readout. Specifically, Vero E6 cells were pre-incubated with each compound dilution for 16 hours, followed by infection with an independent SARS-CoV-2 isolate (SARS-CoV-2 USA-WA1/2020) (MOI = 0.75). 24 hours post-infection, cells were fixed and immunostained for the CoV nucleoprotein (NP). Cellular nuclei were stained with DAPI, prior to automated imaging and analysis. The percentage of infection for each well was calculated as the ratio of infected cells stained with NP antibody, over the total number of cells. Each of these values was normalized to the average of the DMSO control wells in each plate. Twenty-seven percent of compounds (89 compounds) were found to reduce viral replication by at least 40 % at 2.5 µM (data not shown) when averaging data from at least two replicates. These include compounds that were found to belong to enriched target classes (**Figure 2A**), including retinoic acid receptor agonists (LGD-1550, tretinoin, tamibarotene, acitretin, tazarotene, RBAD), the aldose reductase inhibitor AL 3152, benzodiazepine receptor agonists (ZK-93426, zaleplon GR, pagoclone) and antimalarial drugs (AQ-13 and hanfangchin A), as well as the FDA-approved anti-mycobacterial clofazimine.

### Dose response analysis

Although specific for each compound, therapeutic dose ranges are typically expected to track to cellular EC_50_s well below 1 µM concentrations. Therefore, we conducted a dose response analysis to determine the relationship between compound concentration and antiviral activity. Compound concentrations tested ranged from 1.1 nM to 2.5 µM in the immunofluorescence assay described previously. Total cell counts were used to assess compound cytotoxicity (**Figure S4**). In addition to remdesivir, treatment with 17 compounds resulted in discernable dose-dependent antiviral activities, most of which could be segregated based on broad functional, structural, or target-based classes (**Figure 3A**). 10 of these known drugs have EC_50_s > 750 nM or values that could not be exactly extrapolated (**Figures 3A and Figure S5**), while 7 compounds harbored EC_50_s < 500 nM (6 of which reached > 80% inhibiton of the infection at the highest concentration (**Figure 3B-C**)), suggesting that effective antiviral potency could likely be achieved during therapeutic dosing of a COVID-19 patient. Data are also available at <u>reframedb.org</u>.

**Figure 3:**
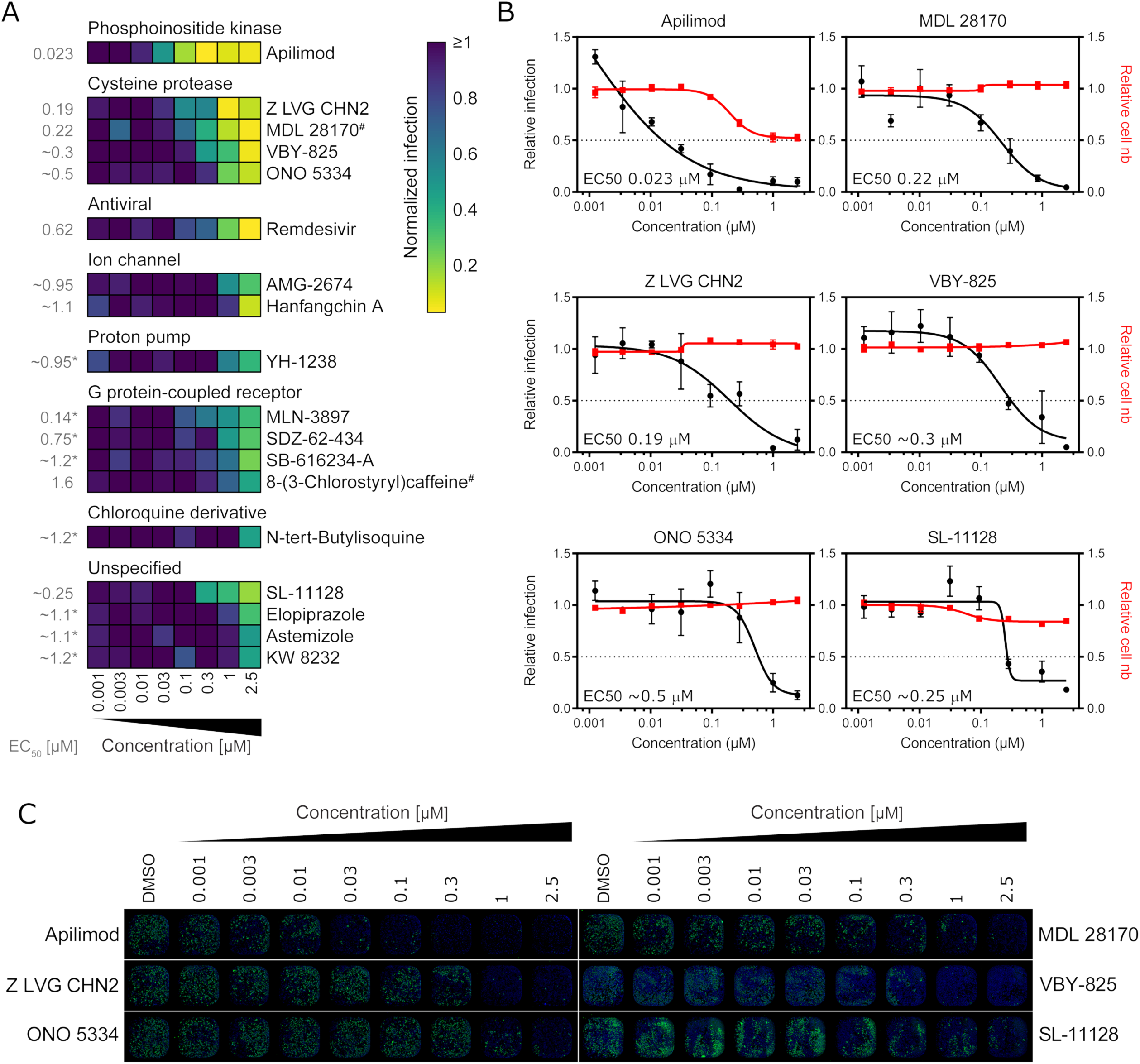
A-C) Vero E6 cells were pre-treated for 16 h with increasing concentrations of the indicated compound and then infected with SARS-CoV-2 at an MOI = 0.75 always in the presence of the compound. 24 h post-infection, cells were fixed and an immunofluorescence was performed, followed by imaging. For each condition, the percentage of infection was calculated as the ratio between the number of infected cells stained for CoV NP and the total amount of cells stained with DAPI. Compound concentrations range between 1 nM and 2.5 µM with 3-fold dilutions. A) Heatmap representing normalized infection of the indicated 17 compounds in dose-response, on a scale from 0 to 1, on the average of three independent experiments. Compounds are associated in clusters, based on their classification category. Concentrations are rounded. # Indicated compounds were evaluated at a concentration of 0.85 µM instead of 1 µM. B) Dose-response curves for both infectivity (black) and cell number (red) are shown. Data are normalized to the average of DMSO-treated wells and represent mean ± SEM for n=3. EC50 for the * indicated compounds was calculated as relative to top concentration. C) Representative immunofluorescence images corresponding to one of the three dose-responses in B are shown. For each condition, the corresponding entire well is shown (4x objective).

To enable prioritization of known drugs for preclinical and clinical evaluation for the treatment of SARS-CoV-2, a summary of the publicly disclosed and relevant preclinical and clinical properties of the most advanced among these molecules are annotated in Table 1 (information retrieved from CortellisTM (Clarivate Analytics) and drugs.com).

**Table 1.**
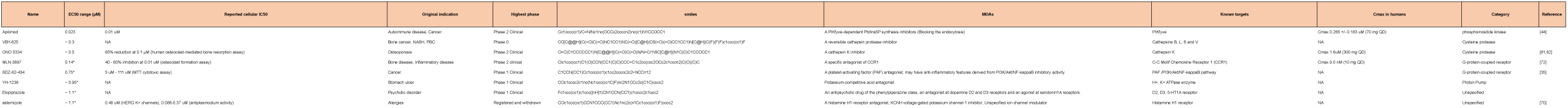
Activity and clinical profiles of compounds with confirmed dose-activity relationships that have entered into clinical evaluation.

## Discussion

Since the beginning of January 2020, an extraordinary number of investigational programs and clinical trials has been initiated in a concerted effort to identify therapeutics against the rapidly growing COVID-19 pandemic. Clinical trials using repurposed clinical-stage or approved drugs such as remdesivir, favipiravir, lopinavir/ritonavir, hydroxychloroquine and others have been under investigation for treating COVID-19 patients^15,16,20,54-59^. Some other therapies, such as treatment with antibodies from seroconverted idividuals, are also being investigated. However, most of the reported studies have been conducted in small cohorts and thus should be considered preliminary, with larger case-control clinical evaluations still pending^15-16,19-22^

The elucidation of additional candidate therapies would greatly enhance the probability of rapidly identifying safe and efficacious treatment options, and would also enable the development of combinatorial regimens (“cocktails”), which reflects the current treatment strategies for HIV-1 and hepatitis C virus (HCV)^60-62^.

Large-scale surveys of existing drugs that may harbor antiviral activities can significantly facilitate such repositioning efforts. A recently reported SARS-CoV-2-human protein-protein interaction (PPI) analysis identified 332 viral-host interactions, 66 of which are potential druggable human host factors targeted by 69 known drugs that have FDA approval, however activities on SARS-CoV-2 replication have not yet been reported^63^. An additional study that tested a focused panel of 48 FDA-approved drugs, previously shown to have antiviral activity against both SARS-CoV and MERS-CoV, also demonstrated that several known drugs harbor potent antiviral activities against SARS-CoV-2^64^.

Here, we report the high-throughput analysis of approximately 12,000 known drugs evaluated for activity against SARS-CoV-2 replication. The assay, conducted in Vero E6 cells was designed to capture multicycle replication, based upon low viral input (MOI = 0.01) and an extended endpoint measurement (72 hours post-infection). Although cell-based assays can be biased towards capturing inhibitors of viral entry, the assay was constructed to interrogate each step of the viral life cycle. Of note, one potential limitation of Vero cells is that, due to species differences, pro-drugs that require the human host cell machinery for processing into their active form, such as some nucleoside inhibitors, may not harbor the same potency as in human cells. Consistently, we found that remdesivir inhibits SARS-CoV-2 replication ∼60-fold more potently in human cells in comparison to Vero E6 cells (**Figure 3A and Figure S5; data not shown**).

The dynamic range of the viral-induced CPE in the assay was small (∼2-2.5 fold), but robust and reproducible (**Figure 1B**). Both the optimization using the LOPAC® 1280library and the first ReFRAME collection screen displayed acceptable Z’ factors (0.4 and 0.51, respectively). The duplicate ReFRAME screen, had a reduced dynamic range (1.5-fold) and corresponding Z’ factor (0.19). Although the correlation between the two ReFRAME replications was high (R^2^=0.68), there were compounds that were found active in replicate 1, but not replicate 2 (**Figure 1C, bottom right of middle panel**). While we leveraged all datasets to select molecules for further validation, compound selection was weighted for replicate 1. Specifically, in addition to 28 molecules from the LOPAC®1280 library, 250 drugs were selected based on their activity. 48 additional ones, belonging to enriched target/MOA sets based on GSEA analysis, were also included.

These selected compounds were tested in an orthogonal assay that directly measures viral replication, in contrast to the indirect measurement of replication assessed by CPE. Here, we took advantage of a high-throughput immunofluorescence assay that monitors infection levels as reflected by SARS-CoV-2 N protein expression at a single cell resolution. Importantly, this validation step enables the separation of molecules that function to block CPE (i.e. cell death), instead of direct effects on replication. This assay was found to be most robust at a 24-hour timepoint using an MOI of 0.75, thus, the antiviral activities of compounds were not interrogated under the original MOI or 72-hour timepoint conditions. Both the earlier timepoint and higher MOI likely biased the validation screen towards the confirmation of early stage inhibitors. Consistent with this hypothesis, we found that several molecules with potent EC_50_s were only able to inhibit replication to approximately 50-60 % at multiple high concentrations, including MLN-3897, YH-1238 and SL-11128 (**Figure 3A and Figure S5**). While this may represent the maximal ability of these molecules to suppress viral replication, alternatively, analysis of these molecules utilizing lower MOIs at later timepoint may reveal greater inhibition of infection. In addition, validation assays were conducted employing drug concentrations that were half of what was utilized in the screen (2.5 μM versus 5 μM) and using a second isolate of SARS-CoV-2. The introduction of these stringencies during the validation step, as well as false positive activities from the HTS assay, likely account for ∼30 % confirmation rate observed at this step of the analysis.

Tretinoin, clofazimine, acitretin were amongst the notable validated compounds, since they have been approved by the FDA. Clofazimine is a lipophilic riminophenazine antibiotic, with described antimycobacterial and anti-inflammatory activity^65^ used for the treatment of leprosy. Main adverse effects include changes in skin pigmentation, nausea and vomiting. The antibacterial activity of clofazimine is described to be related to its ability to bind to the bacterial DNA. Interestingly, this compound was also identified as a potent antiparasitic drug active against *Cryptosporidium*, during a repurposing screen of the ReFRAME library^30^. Further studies are required to understand the mechanism by which this molecule blocks the replication of this positive-strand RNA virus, and determination of the dose-response relationship for clofazimine will enable assessment of whether antiviral efficacy can be achieved at therapeutic doses. Acitretin is an approved orally bioavailable retinoid used for the treatment of psoriasis^66^. Tretinoin, together with tamibarotene, which is registered in Japan, as well as LGD-155, tamibarotene, tazarotene and RBAD, were all validated compounds belonging to the enriched GSEA class of class of retinoic acid agonists highly enriched in the GSEA analysis (**Figure 2A**). It is currently unclear how the activation of the transcriptional program governed by retinoic acid receptors may impinge upon SARS-CoV-2 replication.

Six additional compounds which confirmed in our validation studies also modulate targets that were enriched in the high-throughput screen. Those include aldose reductase inhibitor AL 3151, the benzodiazepine receptor agonists ZK-93426, zaleplon GR and pagoclone, and the two antimalarial drugs AQ-13 and hanfangchin A. Antimalarial drugs have been reported to effectively block several viral infections^67^, including SARS-CoV-2^26^. However, the activities of many of these, in particular chloroquine derivatives, have not been recapitulated in clinical trials^17,18^. Drugs belonging to this class are generally reported to be less effective in blocking viral infections compared to their anti-malarial activity, with EC_50_ generally in a μM range^67^. The concentration required for their antiviral activity could thus likely be difficult to reach in humans, without causing adverse effects. Taken together, confirmation of compounds with membership in enriched target classes underscore the importance of these molecular circuits in the regulation of SARS-CoV-2 replication and support the evaluation of additional preclinical and clinical stage molecules that target over-represented mechanisms.

Among the 17 compounds validated to show a dose-response relationship in our orthogonal assay, 10 compounds harbored EC_50_ antiviral activities >750 nM, suggesting that additional preclinical studies will likely be required to determine whether administration of these compounds can achieve sufficient systemic exposure to enable antiviral activity (**Figure S5**). Seven molecules were found to inhibit viral replication at EC_50_ concentration <500 nM. These include Z LVG CHN2, a preclinical tripeptide derivative that displays a broad-spectrum bactericidal activity. Specifically, this molecule has been previously shown to suppress herpes simplex virus (HSV) replication by inhibiting the enzymatic activity of HSV-encoded cysteine protease^68^, which may indicate that the antiviral function of Z LVG CHN2 occurs through inhibition of SARS-COV-2 3CLpro protease. Another preclinical molecule that exhibits strong antiviral activity, MDL 28170, is a potent cell permeable calpain I and II inhibitor. Interestingly, MDL 28170 was previously found to impair infection by SARS-COV-1 and Ebola virus (EBOV)^50,69^. Additionally, astemizole a registered anti-histamine H1 receptor antagonist that also reported to have anti-malarial properties^70^, inhibited replication at an EC_50_ concentration of ∼1.1 μM. Due to fatal arrhythmias when given in high doses or in combination with certain other common drugs, astemizole has been withdrawn in many countries^71^. Therefore, thorough safety studies are required to determine if there exists a sufficient therapeutic index for the acute treatment of SARS-CoV-2 infection.

MLN-3897 (AVE-9897) is an orally active chemokine CCR1 antagonist and was evaluated in phase II clinical studies for the treatment of rheumatoid arthritis (RA) and multiple sclerosis (MS)^72^, showing a dose of 10 mg once daily was well tolerated^73^. It was determined to inhibit SARS-CoV-2 replication at an estimated EC_50_ concentration of 140 nM (**Figure 2**), and the C_max_ of the compound has been reported at 9.0 nM (10 mg QD). Therefore, additional *in vivo* studies will be required to determine if sufficient systemic concentrations can be reached to promote antiviral activities. The mechanism by which CCR1 antagonism inhibits SARS-CoV-2 infection requires further investigation. However, it has been reported that CCR1 inhibition with MLN-3897 potentially blocks ERK phosphorylation, leading to suppression of the mitogen-activated protein kinase (Raf/MEK/ERK) signal transduction pathway^74^. Interestingly, Raf/MEK/ERK signaling pathways are employed by SARS-CoV-1 to support its replication via multiple well-documented mechanisms^75,76^, and thus this signaling axis may also represent a critical therapeutic target for host-directed SARS-CoV-2 antivirals.

Human cysteinyl cathepsins, including cathepsin B, cathepsin L, and cathepsin K, are required for the proteolytic processing of virally encoded proteins during infection^77-79^. Cathepsin activity seems to be required for proper processing of the SARS-CoV-1 S protein within the endosome in order to activate its fusogenic acitivity^78^. Inhibition of cathepsin L activity has been previously shown to efficiently suppress SARS-CoV-1 infection^78^. We found ONO 5334 (a cathepsin K inhibitor) and VBY-825 (a reversible cathepsin protease inhibitor) to inhibit SARS-CoV-2 infection in a dose-dependent manner, however additional studies will be required to determine if their antiviral activities are due to inhibiting proteolysis of viral or host cellular proteins. ONO 5334 harbored an antiviral EC50 of ∼500 nM, which is in range of a previously reported 85 % activity observed at 100 nM in an osteoclast-mediated bone resorption assay^80^. Importantly, the Cmax of this compound is 1.6 μM (300 mg QD), and treatment with ONO-5334 was well tolerated up to daily doses of 300 mg and for up to 12 months without any clinically relevant safety concerns. ONO5334 reached phase II clinical trials for the treatment of osteoporosis in postmenopausal women, but development was discontinued due to an unfavorable competitive landscape^81,82^. VBY-825, which is in preclinical development, is another cathepsin inhibitor harboring potential antiviral activities against SARS-CoV-2 with an EC50 of ∼300 nM, and it shows high potency against cathepsins B, L, S and V *in vitro*^*83*^. Overall, the identification of VBY-825 and ONO 5334 as effective antiviral molecules against SAR-COV-2 supports the repositioning of these, and potentially additional protease inhibitors, for the treatment of COVID-19 disease.

Finally, apilimod, a specific PIKfyve kinase inhibitor, was found to inhibit SARS-CoV-2 replication at an EC_50_ concentration of 23 nM (**Figure 3**). Importantly, apilimod is found to be well tolerated in humans showing a desirable safety profile at doses of ≤ 125 mg BID, and the Cmax of this compound is 0.265 +/- 0.183 μM (70 mg QD)^84,85^. These data indicate that therapeutic dosing of apilimod in patients can achieve concentrations that are likely to promote antiviral activity. Apilimod has been evaluated in phase II clinical trials for the treatment of active Crohn’s disease and rheumatoid arthritis (RA)^86^, and in additional phase II trials for the oral treatment of common variable immunodeficiency (CVID), but did not show efficacy for these indications^85,87^. In 2019, orphan drug designation was granted to apilimod in the U.S. for the treatment of follicular lymphoma^88^, with phase I clinical trials ongoing. Notably, it has been reported that apilimod efficiently inhibits EBOV, Lassa virus (LASV), and Marburg virus (MARV) in human cell lines, underscoring its potential broad-spectrum antiviral activity^44,89^. The underlying mechanism for the inhibition of SARS-CoV-2 infection by apilimod is currently not known. However, since PIKfyve predominately resides in early endosomes and plays an essential role in maintenance of endomembrane homeostasis, apilimod likely blocks viral low pH-dependent entry through inhibition of the lipid kinase activity of PIKfyve.

Taken together, this study has illuminated a compendium of druggable targets, pathways, biological processes and small molecules that modulate the SARS-CoV-2 replication cycle. These data can be leveraged to focus studies aimed at deciphering the biology of this coronavirus, and to guide additional directed repurposing studies, particularly of existing late-stage preclinical assets. Critically, this campaign led to the identification and validation of 30 molecules with antiviral activities against SARS-CoV-2, including four FDA-approved compounds, one drug registered in Japan, and 11 molecules that have entered clinical trials. The availability of human safety and pharmacological data of these molecules is expected to enable rapid preclinical and clinical assessment of these compounds. However, expedited regulatory review under EUA guidelines also provides a rationale for the development of earlier stage candidate molecules that can be deployed for use during this current pandemic outbreak. It is critical that multiple therapeutic options that demonstrate efficacy against SARS-CoV-2 are available to mitigate potential emergence of drug resistance, as well as enable the evaluation of optimal therapeutic cocktails that are broadly curative for COVID-19 disease.

## Acknowledgments

The following reagent was deposited by the Centers for Disease Control and Prevention and obtained through BEI Resources, NIAID, NIH: SARS-Related Coronavirus 2, Isolate USA-WA1/2020, NR-52281. The authors acknowledge Alan Embry for guidance in securing reagents, Richard Glynne for scientific input and editing, Sunnie Yoh for scientific input, Peter Teriete for technical support, Henrietta Nymark-McMahon for manuscript editing, Tara Marathe for copy editing, Sylvie Blondelle and Larry Adelman for facilities and biosafety support, Marisol Chacon for administrative assistance, Wdee Thienphrapa and Heather Curry for literature research support and Rowland Eaden for shipping and delivery assistance. The authors would also like to thank Olivier Harismendy (University of California, San Diego) for advice on data analysis and Randy Albrecht and Wen-Chun Liu for oversight and administration of the conventional BSL3 facility at the Icahn School of Medicine at Mount Sinai.

This work was supported by the following grants to the Sanford-Burnham-Prebys Medical Discovery Institute: DoD: W81XWH-20-1-0270; DHIPC: U19 AI118610; Fluomics/NOSI: U19 AI135972. The study was partly supported by the donations of Richard Yu and Carol Yu, the Shaw Foundation of Hong Kong, Michael Seak-Kan Tong, and May Tam Mak Mei Yin; and funding from the Theme-Based Research Scheme (T11/707/15) of the Research Grants Council; Hong Kong Special Administrative Region. The work at Calibr at Scripps Research was supported by the Bill & Melinda Gates Foundation.This research was also partly funded by CRIP (Center for Research for Influenza Pathogenesis), a NIAID supported Center of Excellence for Influenza Research and Surveillance (CEIRS, contract # HHSN272201400008C) to AG-S and by supplements to NIAID grant U19AI135972 and DoD grant W81XWH-19-PRMRP-FPA to SKC and AG-S. The funding sources had no role in the study design, data collection, analysis, interpretation, or writing of the report.

---

**Figure S1.**
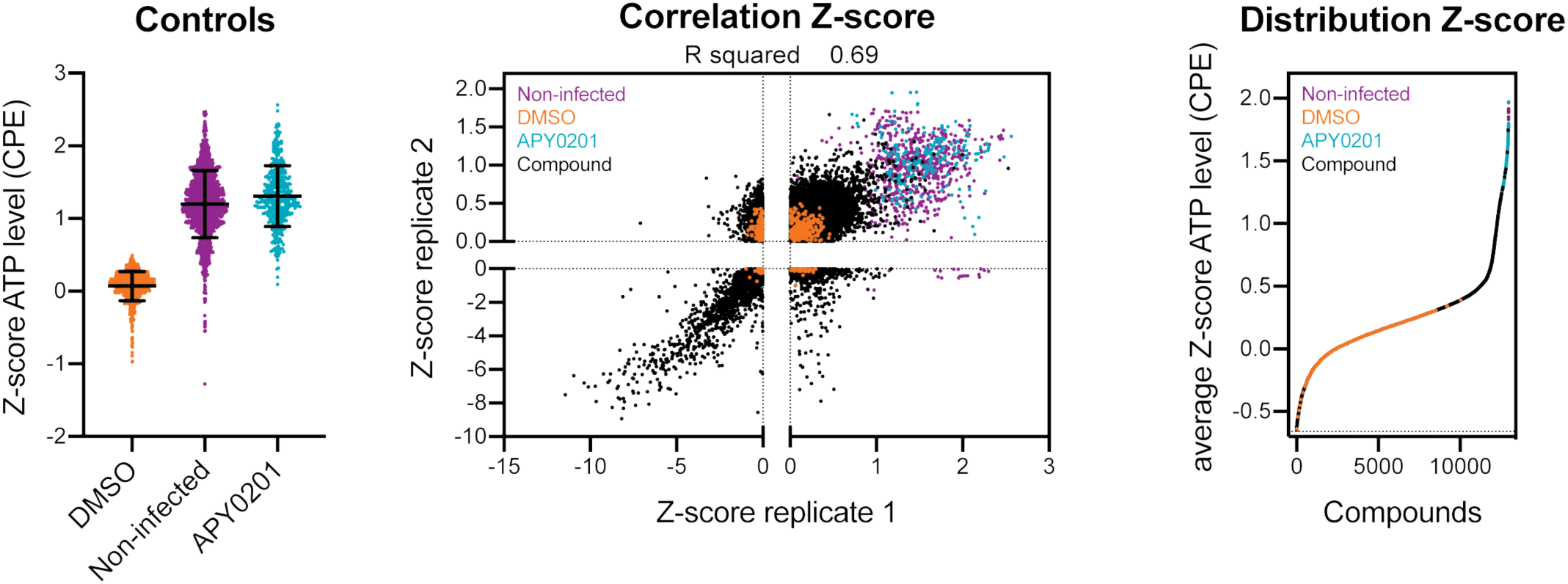
A) Z-scores for ReFRAME collection primary screen. The left graph represents the Z-score of ATP levesl after normalization to the median of each plate for all positive (APY0201) and negative (DMSO) controls as well as for non-infected cells, across all the screening plates. The correlation plot in the middle panel indicates the Z-score of each compound in the two replicates. The distribution of each compound according to the average of the Z-score of each replicate (right panel) is also represented. Each dot indicates the Z-score of each drug in each replicate of the screen (black dots). Values corresponding to DMSO (orange dots), APY0201 (cyan dots) and non-infected cells (purple dots) are also represented. R squared indicates the correlation coefficient for the replicates.

**Figure S2.**
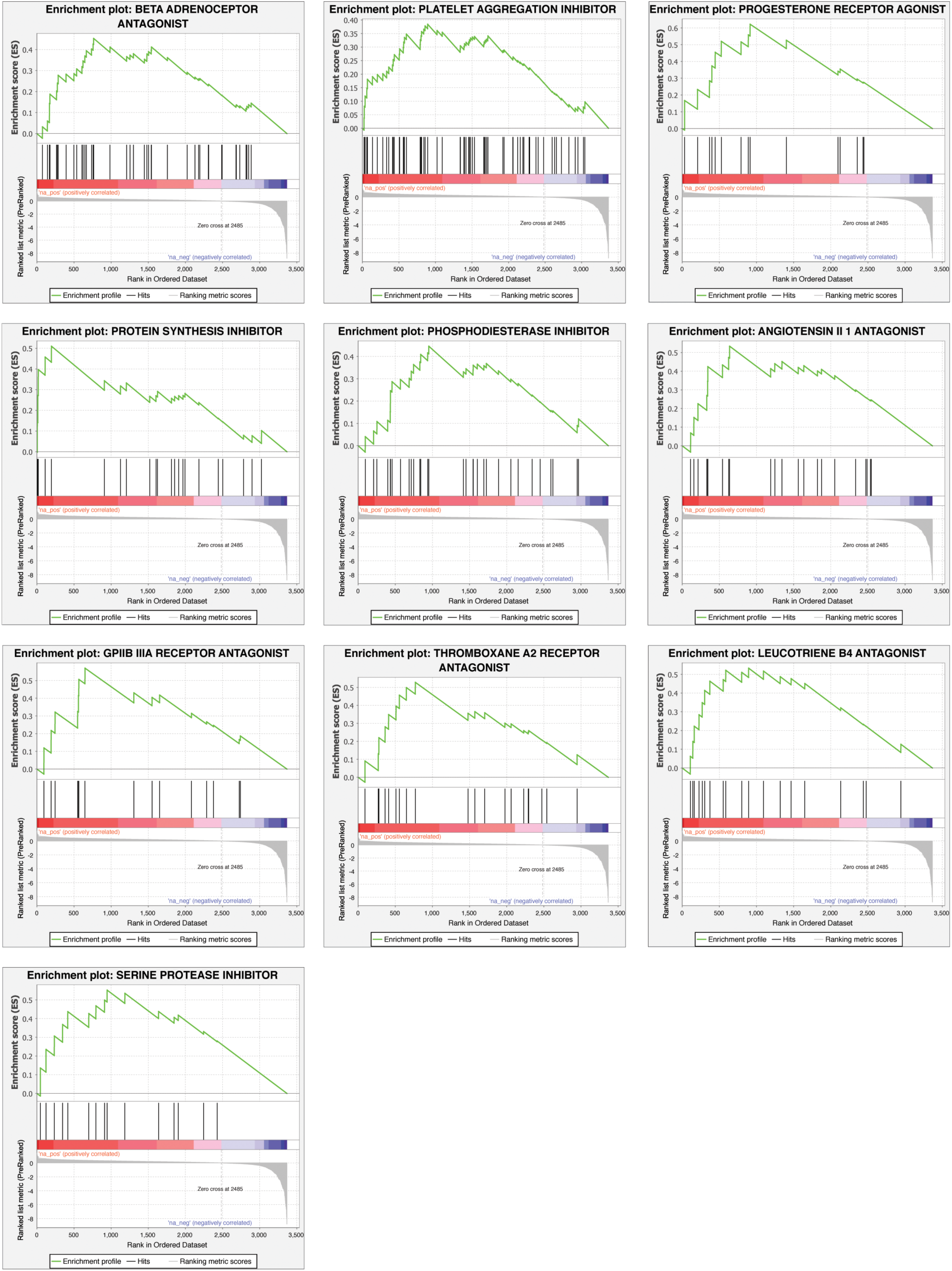
Gene set enrichment analysis (GSEA) of primary screening data according to the average Z′ factor. GSEA enrichment plots of additional ten target clusters that are enriched were represented including beta adrenoreceptor antagonist, platelet aggregation inhibitor, progesterone receptor agonist, protein synthesis inhibitor, phosphodiesterase inhibitor, angiotensin II 1 antagonist, GPIIB IIIA receptor antagonist, thromboxane A2 receptor antagonist, leucotriene B4 antagonist, serine protease inhibitor (P-value < 0.05, FDR q-value < 0.25).

**Figure S3.**
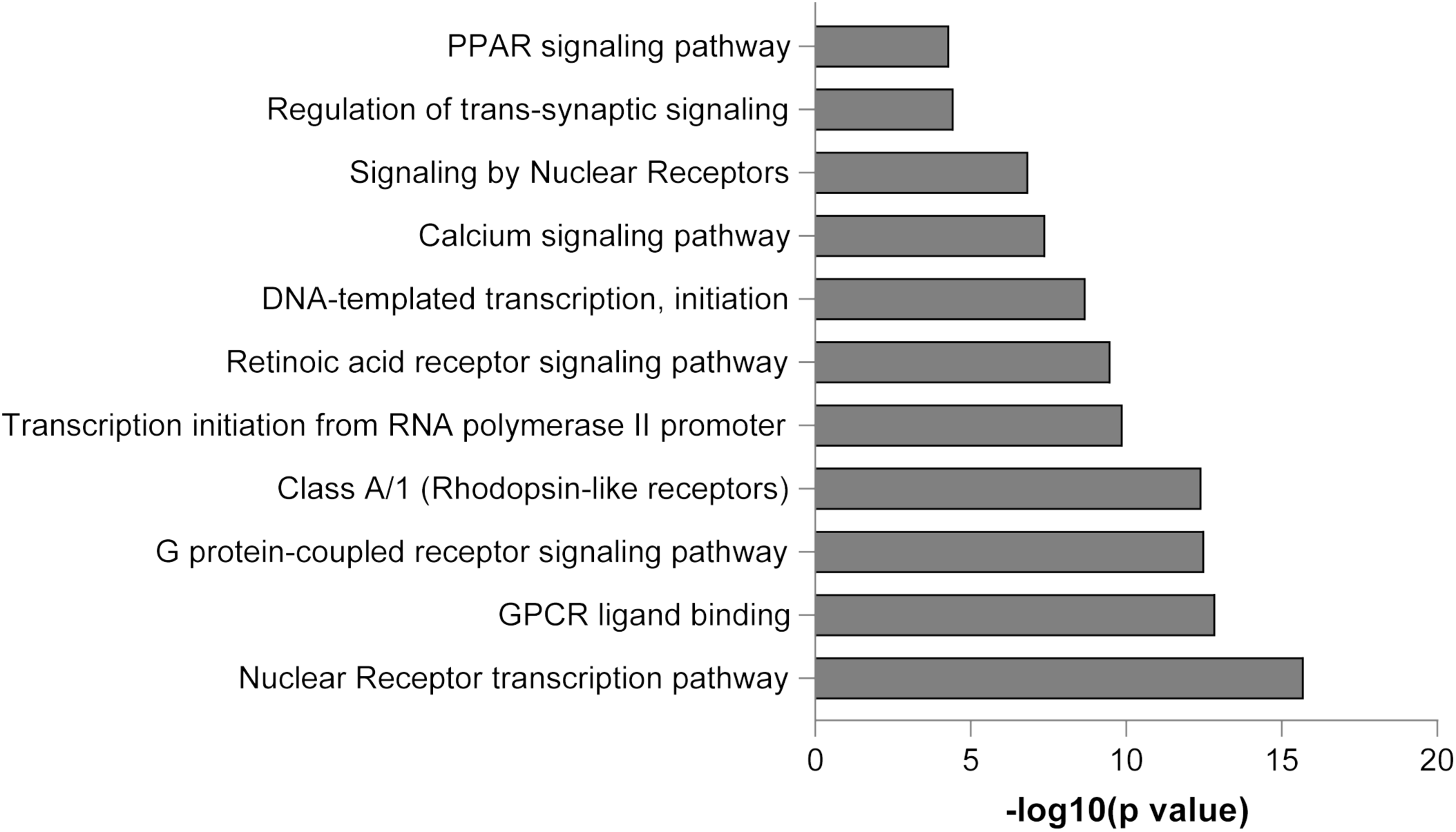
Bar plot of enriched terms across the enriched genes targeted by the compounds. The x-axis corresponds to -log10(p value) while the y-axis indicates the enriched terms. The analysis was performed using Metascape^40^.

**Figure S4.**
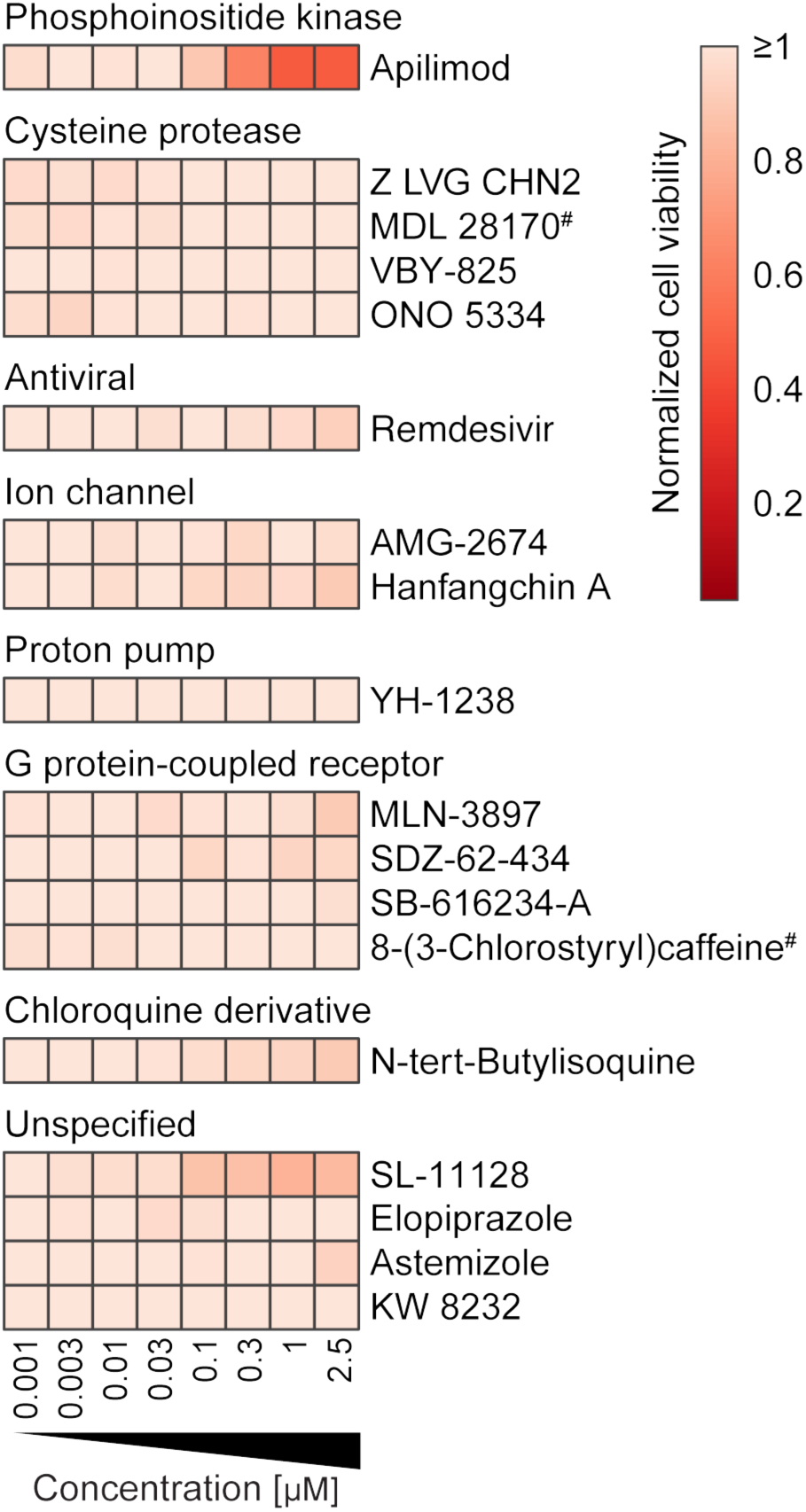
Vero E6 cells were pre-treated for 16 h with increasing concentrations of the indicated compound and then infected with SARS-CoV-2 at an MOI = 0.75 always in the presence of the compound. 24 h post-infection, cells were fixed and an immunofluorescence was performed, followed by imaging. For each condition, the total amount of cells stained with DAPI was calculated. Data are normalized to the average of DMSO-treated wells. The heatmap represents the normalized cell number of the indicated 17 compounds in dose-response, on a scale from 0 to 1, on the average of three independent experiments. Compounds are associated in clusters, based on their classification category. Concentrations are rounded. # Indicated compounds were evaluated at a concentration of 0.85 µM instead of 1 µM.

**Figure S5.**
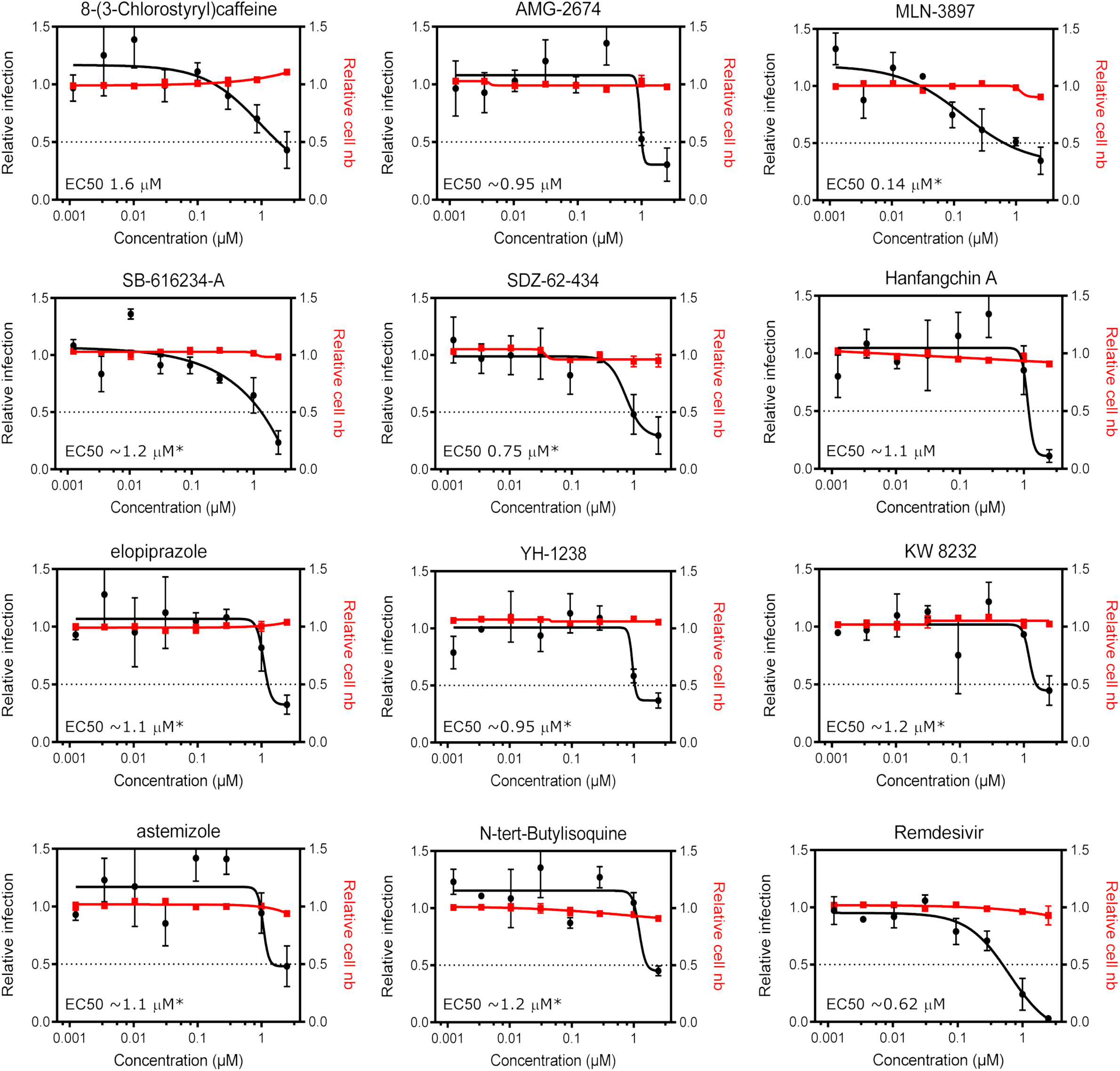
Vero E6 cells were pre-treated for 16 h with increasing concentrations of the indicated compound and then infected with SARS-CoV-2 at an MOI = 0.75 always in the presence of the compound. 24 h post-infection, cells were fixed and an immunofluorescence was performed. For each condition, the percentage of infection was calculated as the ratio between the number of infected cells stained for CoV NP and the total amount of cells stained with DAPI. Dose-response curves for both infectivity (black) and cell number (red) are shown. Data are normalized to the average of DMSO-treated wells and represent mean ± SEM for n=3. EC50 for the * indicated compounds was calculated as relative to top concentration.

**Table S1**

Drug targets enriched in GSEA analysis of HTS data.

**Table S2**

FDA-approved or GSEA-enriched compounds.

## Notes

### Competing Interest Statement

The authors have declared no competing interest.

## References

1 Yang, X. et al. Clinical course and outcomes of critically ill patients with SARS-CoV-2 pneumonia in Wuhan, China: a single-centered, retrospective, observational study. Lancet Respir Med, doi: 10.1016/S2213-2600(20)30079-5 (2020).

2 World_Health_Organization. https://www.who.int/emergencies/diseases/novel-coronavirus-2019, 2020).

3 World_Health_Organization. (2020).

4 Onder, G., Rezza, G. & Brusaferro, S. Case-Fatality Rate and Characteristics of Patients Dying in Relation to COVID-19 in Italy. JAMA, doi: 10.1001/jama.2020.4683 (2020).

5 Jin, X. et al. Epidemiological, clinical and virological characteristics of 74 cases of coronavirus-infected disease 2019 (COVID-19) with gastrointestinal symptoms. Gut, doi: 10.1136/gutjnl-2020-320926 (2020).

6 Rodriguez-Morales, A. J. et al. Clinical, laboratory and imaging features of COVID-19: A systematic review and meta-analysis. Travel Med Infect Dis, 101623, doi: 10.1016/j.tmaid.2020.101623 (2020).

7 ClincalTrials.Gov. https://clinicaltrials.gov/ct2/results?cond=COVID-19&term=sarilumab&cntry=&state=&city=&dist=&Search=Search.

8 Center_For_Disease_Control. https://www.cdc.gov/coronavirus/2019-ncov/need-extra-precautions/people-at-higher-risk.html. (2020).

9 Livingston, E. & Bucher, K. Coronavirus Disease 2019 (COVID-19) in Italy. JAMA, doi: 10.1001/jama.2020.4344 (2020).

10 Remuzzi, A. & Remuzzi, G. COVID-19 and Italy: what next? Lancet, doi: 10.1016/S0140-6736(20)30627-9 (2020).

11 Spinelli, A. & Pellino, G. COVID-19 pandemic: perspectives on an unfolding crisis. Br J Surg, doi: 10.1002/bjs.11627 (2020).

12 Food_And_Drug_Administration. EUA Hydroxychloroquine Phosphate Patients and Parent/Caregivers Fact Sheet, version date 3/28/20.

13 Chen, W. H., Strych, U., Hotez, P. J. & Bottazzi, M. E. The SARS-CoV-2 Vaccine Pipeline: an Overview. Current tropical medicine reports, 1–4, doi: 10.1007/s40475-020-00201-6 (2020).

14 Cascella, M., Rajnik, M., Cuomo, A., Dulebohn, S. C. & Di Napoli, R. in StatPearls (StatPearls Publishing StatPearls Publishing LLC., 2020).

15 Gao, J., Tian, Z. & Yang, X. Breakthrough: Chloroquine phosphate has shown apparent efficacy in treatment of COVID-19 associated pneumonia in clinical studies. Biosci Trends 14, 72–73, doi: 10.5582/bst.2020.01047 (2020).

16 Gautret, P. et al. Hydroxychloroquine and Azithromycin as a treatment of COVID-19: preliminary results of an open-label non-randomized clinical trial. medRxiv, 2020.2003.2016.20037135, doi: 10.1101/2020.03.16.20037135 (2020).

17 Inciardi, R. M. et al. Cardiac Involvement in a Patient With Coronavirus Disease 2019 (COVID-19). JAMA cardiology, doi: 10.1001/jamacardio.2020.1096 (2020).

18 Kapoor, A. et al. Cardiovascular risks of hydroxychloroquine in treatment and prophylaxis of COVID-19 patients: A scientific statement from the Indian Heart Rhythm Society. Indian Pacing and Electrophysiology Journal, doi: https://doi.org/10.1016/j.ipej.2020.04.003 (2020).

19 ClinicalTrials.gov. https://clinicaltrials.gov/ct2/results?cond=COVID-19&term=kaletra&cntry=&state=&city=&dist=&Search=Search.

20 Chen, H. et al. First Clinical Study Using HCV Protease Inhibitor Danoprevir to Treat Naive and Experienced COVID-19 Patients. medRxiv, 2020.2003.2022.20034041, doi: 10.1101/2020.03.22.20034041 (2020).

21 ClinicalTrials.gov. https://clinicaltrials.gov/ct2/results?cond=COVID-19&term=favipiravir&cntry=&state=&city=&dist=&Search=Search.

22 ClinicalTrials.Gov. https://clinicaltrials.gov/ct2/results?cond=COVID-19&term=remdesivir&cntry=&state=&city=&dist=&Search=Search.

23 Warren, T. K. et al. Therapeutic efficacy of the small molecule GS-5734 against Ebola virus in rhesus monkeys. Nature 531, 381–385, doi: 10.1038/nature17180 (2016).

24 Agostini, M. L. et al. Coronavirus Susceptibility to the Antiviral Remdesivir (GS-5734) Is Mediated by the Viral Polymerase and the Proofreading Exoribonuclease. mBio 9, e00221–00218, doi: 10.1128/mBio.00221-18 (2018).

25 Sheahan, T. P. et al. Comparative therapeutic efficacy of remdesivir and combination lopinavir, ritonavir, and interferon beta against MERS-CoV. Nature Communications 11, 222, doi: 10.1038/s41467-019-13940-6 (2020).

26 Wang, M. et al. Remdesivir and chloroquine effectively inhibit the recently emerged novel coronavirus (2019-nCoV) in vitro. Cell Res 30, 269–271, doi: 10.1038/s41422-020-0282-0 (2020).

27 Janes, J. et al. The ReFRAME library as a comprehensive drug repurposing library and its application to the treatment of cryptosporidiosis. Proc Natl Acad Sci U S A 115, 10750–10755, doi: 10.1073/pnas.1810137115 (2018).

28 Kim, Y. J. et al. The ReFRAME library as a comprehensive drug repurposing library to identify mammarenavirus inhibitors. Antiviral Res 169, 104558, doi: 10.1016/j.antiviral.2019.104558 (2019).

29 Harbut, M. B. et al. Auranofin exerts broad-spectrum bactericidal activities by targeting thiol-redox homeostasis. Proc Natl Acad Sci U S A 112, 4453–4458, doi: 10.1073/pnas.1504022112 (2015).

30 Love, M. S. et al. A high-throughput phenotypic screen identifies clofazimine as a potential treatment for cryptosporidiosis. PLoS Negl Trop Dis 11, e0005373, doi: 10.1371/journal.pntd.0005373 (2017).

31 Bollong, M. J. et al. Small molecule-mediated inhibition of myofibroblast transdifferentiation for the treatment of fibrosis. Proc Natl Acad Sci U S A 114, 4679–4684, doi: 10.1073/pnas.1702750114 (2017).

32 Corsello, S. M. et al. The Drug Repurposing Hub: a next-generation drug library and information resource. Nat Med 23, 405–408, doi: 10.1038/nm.4306 (2017).

33 Li, Y. Y. & Jones, S. J. Drug repositioning for personalized medicine. Genome Med 4, 27, doi: 10.1186/gm326 (2012).

34 Ashburn, T. T. & Thor, K. B. Drug repositioning: identifying and developing new uses for existing drugs. Nat Rev Drug Discov 3, 673–683, doi: 10.1038/nrd1468 (2004).

35 Brunton, V. G. & Workman, P. In vitro antitumour activity of the novel imidazoisoquinoline SDZ 62-434. Br J Cancer 67, 989–995, doi: 10.1038/bjc.1993.181 (1993).

36 To, K. K. et al. Consistent detection of 2019 novel coronavirus in saliva. Clin Infect Dis, doi: 10.1093/cid/ciaa149 (2020).

37 Yuan, S. et al. SREBP-dependent lipidomic reprogramming as a broad-spectrum antiviral target. Nat Commun 10, 120, doi: 10.1038/s41467-018-08015-x (2019).

38 Subramanian, A. et al. Gene set enrichment analysis: A knowledge-based approach for interpreting genome-wide expression profiles. Proceedings of the National Academy of Sciences 102, 15545–15550, doi: 10.1073/pnas.0506580102 (2005).

39 Mootha, V. K. et al. PGC-1α-responsive genes involved in oxidative phosphorylation are coordinately downregulated in human diabetes. Nature Genetics 34, 267–273, doi: 10.1038/ng1180 (2003).

40 Zhou, Y. et al. Metascape provides a biologist-oriented resource for the analysis of systems-level datasets. Nat Commun 10, 1523, doi: 10.1038/s41467-019-09234-6 (2019).

41 Deprez, M. et al. A single-cell atlas of the human healthy airways. bioRxiv, 2019.2012.2021.884759, doi: 10.1101/2019.12.21.884759 (2019).

42 Matsuyama, S. et al. Enhanced isolation of SARS-CoV-2 by TMPRSS2-expressing cells. Proc Natl Acad Sci U S A 117, 7001–7003, doi: 10.1073/pnas.2002589117 (2020).

43 Park, W. B. et al. Virus Isolation from the First Patient with SARS-CoV-2 in Korea. J Korean Med Sci 35, e84, doi: 10.3346/jkms.2020.35.e84 (2020).

44 Nelson, E. A. et al. The phosphatidylinositol-3-phosphate 5-kinase inhibitor apilimod blocks filoviral entry and infection. PLoS Negl Trop Dis 11, e0005540, doi: 10.1371/journal.pntd.0005540 (2017).

45 Qiu, S. et al. Ebola virus requires phosphatidylinositol (3,5) bisphosphate production for efficient viral entry. Virology 513, 17–28, doi: 10.1016/j.virol.2017.09.028 (2018).

46 Yamamoto, N. et al. HIV protease inhibitor nelfinavir inhibits replication of SARS-associated coronavirus. Biochemical and biophysical research communications 318, 719–725, doi: 10.1016/j.bbrc.2004.04.083 (2004).

47 Shweta, C., Yashpal S., M. & Shailly, T. Identification of SARS-CoV-2 Cell Entry Inhibitors by Drug Repurposing Using in Silico Structure-Based Virtual Screening Approach. (2020).

48 Zhijian, X. et al. Nelfinavir Is Active Against SARS-CoV-2 in Vero E6 Cells. (2020).

49 Wang, S. Q. et al. Virtual screening for finding natural inhibitor against cathepsin-L for SARS therapy. Amino Acids 33, 129–135, doi: 10.1007/s00726-006-0403-1 (2007).

50 Schneider, M. et al. Severe acute respiratory syndrome coronavirus replication is severely impaired by MG132 due to proteasome-independent inhibition of M-calpain. J Virol 86, 10112–10122, doi: 10.1128/jvi.01001-12 (2012).

51 Mootha, V. K. et al. PGC-1alpha-responsive genes involved in oxidative phosphorylation are coordinately downregulated in human diabetes. Nat Genet 34, 267–273, doi: 10.1038/ng1180 (2003).

52 Subramanian, A. et al. Gene set enrichment analysis: a knowledge-based approach for interpreting genome-wide expression profiles. Proc Natl Acad Sci U S A 102, 15545–15550, doi: 10.1073/pnas.0506580102 (2005).

53 Lukassen, S. et al. SARS-CoV-2 receptor ACE2 and TMPRSS2 are primarily expressed in bronchial transient secretory cells. The EMBO Journal n/a, e105114, doi: 10.15252/embj.2020105114 (2020).

54 Bian, H. et al. Meplazumab treats COVID-19 pneumonia: an open-labelled, concurrent controlled add-on clinical trial. medRxiv, 2020.2003.2021.20040691, doi: 10.1101/2020.03.21.20040691 (2020).

55 Cao, B. et al. A Trial of Lopinavir-Ritonavir in Adults Hospitalized with Severe Covid-19. N Engl J Med, doi: 10.1056/NEJMoa2001282 (2020).

56 Kujawski, S. A. et al. First 12 patients with coronavirus disease 2019 (COVID-19) in the United States. medRxiv, 2020.2003.2009.20032896, doi: 10.1101/2020.03.09.20032896 (2020).

57 Marmor, M. F. et al. Recommendations on Screening for Chloroquine and Hydroxychloroquine Retinopathy (2016 Revision). Ophthalmology 123, 1386–1394, doi: 10.1016/j.ophtha.2016.01.058 (2016).

58 Wang, Z., Chen, X., Lu, Y., Chen, F. & Zhang, W. Clinical characteristics and therapeutic procedure for four cases with 2019 novel coronavirus pneumonia receiving combined Chinese and Western medicine treatment. Biosci Trends 14, 64–68, doi: 10.5582/bst.2020.01030 (2020).

59 Zhang, Q., Wang, Y., Qi, C., Shen, L. & Li, J. Clinical trial analysis of 2019-nCoV therapy registered in China. J Med Virol, doi: 10.1002/jmv.25733 (2020).

60 Matthew, A. N., Kurt Yilmaz, N. & Schiffer, C. A. Mavyret: A Pan-Genotypic Combination Therapy for the Treatment of Hepatitis C InfectionPublished as part of the Biochemistry series “Biochemistry to Bedside”. Biochemistry 57, 481–482, doi: 10.1021/acs.biochem.7b01160 (2018).

61 Ferenci, P. New anti-HCV drug combinations: who will benefit? The Lancet. Infectious diseases 17, 1008–1009, doi: 10.1016/S1473-3099(17)30486-3 (2017).

62 Cihlar, T. & Fordyce, M. Current status and prospects of HIV treatment. Current opinion in virology 18, 50–56, doi: 10.1016/j.coviro.2016.03.004 (2016).

63 Gordon, D. E. et al. A SARS-CoV-2-Human Protein-Protein Interaction Map Reveals Drug Targets and Potential Drug-Repurposing. bioRxiv, 2020.2003.2022.002386, doi: 10.1101/2020.03.22.002386 (2020).

64 Weston, S., Haupt, R., Logue, J., Matthews, K. & Frieman, M. B. FDA approved drugs with broad anti-coronaviral activity inhibit SARS-CoV-2 <em>in vitro</em>. bioRxiv, 2020.2003.2025.008482, doi: 10.1101/2020.03.25.008482 (2020).

65 Garrelts, J. C. Clofazimine: a review of its use in leprosy and Mycobacterium avium complex infection. DICP 25, 525–531, doi: 10.1177/106002809102500513 (1991).

66 Lee, C. S. & Li, K. A review of acitretin for the treatment of psoriasis. Expert Opin Drug Saf 8, 769–779, doi: 10.1517/14740330903393732 (2009).

67 D’Alessandro, S. et al. The Use of Antimalarial Drugs against Viral Infection. Microorganisms 8, doi: 10.3390/microorganisms8010085 (2020).

68 Bjorck, L., Grubb, A. & Kjellen, L. Cystatin C, a human proteinase inhibitor, blocks replication of herpes simplex virus. J Virol 64, 941–943 (1990).

69 Zhou, Y. & Simmons, G. Development of novel entry inhibitors targeting emerging viruses. Expert Rev Anti Infect Ther 10, 1129–1138, doi: 10.1586/eri.12.104 (2012).

70 Chong, C. R., Chen, X., Shi, L., Liu, J. O. & Sullivan, D. J., Jr. A clinical drug library screen identifies astemizole as an antimalarial agent. Nature chemical biology 2, 415–416, doi: 10.1038/nchembio806 (2006).

71 de Abajo, F. J. & Rodriguez, L. A. Risk of ventricular arrhythmias associated with nonsedating antihistamine drugs. Br J Clin Pharmacol 47, 307–313, doi: 10.1046/j.1365-2125.1999.00885.x (1999).

72 Vallet, S. et al. MLN3897, a novel CCR1 inhibitor, impairs osteoclastogenesis and inhibits the interaction of multiple myeloma cells and osteoclasts. Blood 110, 3744–3752, doi: 10.1182/blood-2007-05-093294 (2007).

73 Vergunst, C. E. et al. MLN3897 plus methotrexate in patients with rheumatoid arthritis: safety, efficacy, pharmacokinetics, and pharmacodynamics of an oral CCR1 antagonist in a phase IIa, double-blind, placebo-controlled, randomized, proof-of-concept study. Arthritis and rheumatism 60, 3572–3581, doi: 10.1002/art.24978 (2009).

74 Vallet, S. et al. A novel role for CCL3 (MIP-1α) in myeloma-induced bone disease via osteocalcin downregulation and inhibition of osteoblast function. Leukemia 25, 1174–1181, doi: 10.1038/leu.2011.43 (2011).

75 Cai, Y., Liu, Y. & Zhang, X. Suppression of Coronavirus Replication by Inhibition of the MEK Signaling Pathway. Journal of Virology 81, 446–456, doi: 10.1128/jvi.01705-06 (2007).

76 Pleschka, S. RNA viruses and the mitogenic Raf/MEK/ERK signal transduction cascade. Biological chemistry 389, 1273–1282, doi: 10.1515/bc.2008.145 (2008).

77 Mori, Y. et al. Processing of Capsid Protein by Cathepsin L Plays a Crucial Role in Replication of Japanese Encephalitis Virus in Neural and Macrophage Cells. Journal of Virology 81, 8477–8487, doi: 10.1128/jvi.00477-07 (2007).

78 Simmons, G. et al. Inhibitors of cathepsin L prevent severe acute respiratory syndrome coronavirus entry. Proceedings of the National Academy of Sciences of the United States of America 102, 11876–11881, doi: 10.1073/pnas.0505577102 (2005).

79 Marzi, A., Reinheckel, T. & Feldmann, H. Cathepsin B & L Are Not Required for Ebola Virus Replication. PLOS Neglected Tropical Diseases 6, e1923, doi: 10.1371/journal.pntd.0001923 (2012).

80 Duong, L. T. Therapeutic inhibition of cathepsin K-reducing bone resorption while maintaining bone formation. Bonekey Rep 1, 67–67, doi: 10.1038/bonekey.2012.67 (2012).

81 Eastell, R. et al. Safety and efficacy of the cathepsin K inhibitor ONO-5334 in postmenopausal osteoporosis: the OCEAN study. Journal of bone and mineral research : the official journal of the American Society for Bone and Mineral Research 26, 1303–1312, doi: 10.1002/jbmr.341 (2011).

82 Eastell, R. et al. Effect of ONO-5334 on bone mineral density and biochemical markers of bone turnover in postmenopausal osteoporosis: 2-year results from the OCEAN study. Journal of bone and mineral research : the official journal of the American Society for Bone and Mineral Research 29, 458–466, doi: 10.1002/jbmr.2047 (2014).

83 Elie, B. T. et al. Identification and pre-clinical testing of a reversible cathepsin protease inhibitor reveals anti-tumor efficacy in a pancreatic cancer model. Biochimie 92, 1618–1624, doi: 10.1016/j.biochi.2010.04.023 (2010).

84 Gayle, S. et al. Identification of apilimod as a first-in-class PIKfyve kinase inhibitor for treatment of B-cell non-Hodgkin lymphoma. Blood 129, 1768–1778, doi: 10.1182/blood-2016-09-736892 (2017).

85 Sbrissa, D., Naisan, G., Ikonomov, O. C. & Shisheva, A. Apilimod, a candidate anticancer therapeutic, arrests not only PtdIns(3,5)P2 but also PtdIns5P synthesis by PIKfyve and induces bafilomycin A1-reversible aberrant endomembrane dilation. PloS one 13, e0204532–e0204532, doi: 10.1371/journal.pone.0204532 (2018).

86 Sands, B. E. et al. Randomized, double-blind, placebo-controlled trial of the oral interleukin-12/23 inhibitor apilimod mesylate for treatment of active Crohn’s disease. Inflammatory bowel diseases 16, 1209–1218, doi: 10.1002/ibd.21159 (2010).

87 Billich, A. Drug evaluation: apilimod, an oral IL-12/IL-23 inhibitor for the treatment of autoimmune diseases and common variable immunodeficiency. IDrugs : the investigational drugs journal 10, 53–59 (2007).

88 Sultana, F. et al. Snx10 and PIKfyve are required for lysosome formation in osteoclasts. Journal of cellular biochemistry 121, 2927–2937, doi: 10.1002/jcb.29534 (2020).

89 Qiu, S. et al. Ebola virus requires phosphatidylinositol (3,5) bisphosphate production for efficient viral entry. Virology 513, 17–28, doi: https://doi.org/10.1016/j.virol.2017.09.028 (2018).

90 Ziegler, C. et al. SARS-CoV-2 Receptor ACE2 is an Interferon-Stimulated Gene in Human Airway Epithelial Cells and Is Enriched in Specific Cell Subsets Across Tissues. CELL-D-20-00767 (2020), doi: http://dx.doi.org/10.2139/ssrn.3555145

